# Importance of a heat snap in RT-PCR quantification of rotavirus double-stranded RNA in wastewater

**DOI:** 10.1101/2025.08.18.670621

**Authors:** Seju Kang, Anna Wettlauffer, Jolinda de Korne-Elenbaas, Charles B. Niwagaba, Linda Strande, Dorothea Duong, Bridgette Shelden, Timothy R. Julian, Alexandria B. Boehm

## Abstract

Quantification of copies of double stranded RNA using RT-PCR methods may require denaturation of the double stranded structure using an initial high temperature incubation followed by rapid cooling, herein called “heat snap”. Papers in the literature that report rotavirus RNA concentrations in fecal and environmental samples do not consistently report the use of such a “heat snap”. In this study, we quantified rotavirus RNA in diverse environmental samples (wastewater solids, wastewater, and drainage samples) using digital RT-PCR methods with and without a heatsnap. Concentrations were higher in samples by a factor of 125 when a heat snap was applied. This was consistent across sample types, and across laboratories and PCR instrumentation. We recommend a heat snap be used when enumerating double stranded RNA from rotavirus and other viruses in environmental samples.

## Introduction

Enumeration of copies of genomic RNA that is natively double stranded is uncommon in molecular biology as dsRNA genomes are amongst the rarest found in viruses; an estimated 3% (430 of 16215) of known viruses have dsRNA genomes^1^. When using a quantitative RT-PCR approach to quantify dsRNA, an initial high temperature incubation step may be needed to denature dsRNA to ensure the RNA template is available for the reverse transcription (RT) enzyme to synthesize cDNA. This heating step is typically accomplished by heating the reaction at 95-100°C for 5 min, and then cooling on ice or at 4°C prior^2^ to RT. Herein we refer to this as a “heat snap”. The heat snap step is sometimes conducted on the extracted dsRNA alone^3^, and sometimes on the dsRNA combined with primers and dNTP ^4^.

Despite the published use of the heat snap to denature dsRNA prior to RT, we find that it is not universally implemented in studies quantifying rotavirus from clinical or environmental samples. In 2024, Awere-Duodu and Donkor^5^ published a systematic review of prevalence rates of rotavirus detection in diverse water environments. The review identified 75 publications that reported detection or quantification of rotavirus in drinking water, ground water, sewage, and surface waters, of which 74 used PCR-based detection of RNA. Amongst these 74 publications, only 17 (23%) reported the steps for the heat snap directly in the methods. An additional 10 (14%) did not include the heat snap steps in the methods, but referenced prior studies that describe clearly the heat snap. The remaining 47 (64%) did not describe a heat snap step or include a reference to a prior study or protocol describing a heat snap step, suggesting the authors did not include or were not aware of this step (compiled data provided at Stanford Digital Repository: https://doi.org/10.25740/rk700qq6039). In many of the studies that did not include a heat snap, reverse transcription kits were referenced that do not include guidance on denaturing dsRNA prior to reverse transcription. Given the substantial diversity in sample types, locations, and extraction and detection methods, we did not attempt to determine the impact of heat snap on prevalence of rotavirus amongst these studies.

We recently published a systematic review of the literature to identify studies that report concentrations of rotavirus RNA in human excretions (for example, feces, sputum, urine)^6^. This set of compiled papers offered a convenient set to examine whether or not researchers report using a heat snap during rotavirus dsRNA quantification from clinical samples. We identified 25 papers that report rotavirus RNA in excretions as measured using quantitative RT-PCR methods, these papers all happened to be focused on RNA quantification in stool. Of these papers, ten ^7–16^ did not report using a heat snap, twelve^17–27^ did report using a heat snap, and three ^28–30^ did not provide cycling conditions or clear references to the cycling conditions and therefore it was not possible to discern whether they used a heat snap. It is not necessarily appropriate to compare measurements obtained in the different studies because they each report concentrations of rotavirus RNA in excretions from individuals experiencing different stages and severities of rotavirus infections when RNA concentrations might be higher or lower due to various biological factors. Despite this limitation, we found that rotavirus RNA concentrations reported in papers describing using a heat snap had significantly higher log_10_ concentrations than those that did not (log_10_-mean of samples with heat snap = 6.9, log_10_-mean without heat snap = 6.3 log_10_copies/g; t-test, t = 4.428, dof = 718, p<10^−4^, compiled data provided at the Stanford Digital Repository: https://doi.org/10.25740/rk700qq6039). These findings suggest not using a heat snap may underestimate the concentration of a dsRNA target, especially if the dsRNA is not already denatured in the media where it is being quantified.

Wastewater measurements of infectious disease biomarkers can allow public health professionals and infectious disease researchers to infer information on the occurrence of diseases in contributing populations^31^. As rotavirus is one of the most important etiologies of diarrheal disease and can cause significant morbidity and mortality in children^32^, it is a potentially important target for wastewater surveillance. Rotavirus RNA has been enumerated in wastewater matrices previously^33–39^, but none of the studies describe using a heat snap in their methods. Rotavirus RNA in wastewater may be encapsulated in damaged or intact viral capsids, or exist external to a capsid, and it may be fragmented, or denatured. If the dsRNA is not denatured in wastewater, then the use of a heat snap may be needed for sensitive and accurate rotavirus RNA quantification.

The goal of the present study was to determine whether inclusion of a heat snap affects quantification of rotavirus RNA in wastewater using digital RT-PCR. The work was carried out in two different laboratories using different instruments.

## Methods

### Quantification of rotavirus RNA in wastewater solids

Wastewater solids were collected from 31 wastewater treatment plants located in 13 states across the United States (Table 1) between 29 April and 27 June 2025. Several plants provided more than one sample yielding a total of n = 38 samples. Samples were provided to Stanford University (Stanford, California, USA) for the WastewaterSCAN project. The methods for wastewater solids sample collection, nucleic-acid extraction are outlined in detail by Boehm et al.^40^ and in published protocols^41,42^ and not repeated herein. Nucleic-acids were stored at −80°C for approximately 7 days prior to analysis. Samples were thawed at 4°C and immediately used as template in droplet digital one-step RT-PCR reactions using rotavirus primers and probes described previously ^43^, labeled using ATTO590. The assay was run in multiplex with previously described assays for parvovirus B19^44^, adenovirus group F^43^, and measles^45^ RNA, the results of those assays are not described herein. The droplet generator, thermocycler, and droplet readers were purchased from Biorad, as described previously^40^. Two different cycling conditions were used. One set of conditions did not include a heat snap and are reported by Boehm et al.^40^ The other set of conditions included an initial denaturation step where the reaction was heated to 99°C for 5 min followed by cooling at 4°C for 5 min. These conditions are subsequently referred to as conditions without and with a heat snap, respectively. Nucleic-acid templates were run in 6 replicate wells and results from the replicate wells were merged for post processing following methods previously described^40^. Positive and negative controls were run as described elsewhere^40^. Concentrations are provided in units of copies per gram dry weight and errors are provided as standard deviations. The lowest detectable concentration was approximately 1000 copies per g dry weight solids.

**Table 1.** Locations (city, state) where wastewater solids were collected in the United States. Number of samples per sit is 1 per site except for Kansas City, KS (n=6), Merced, CA (n=2), and Turlock, CA (n=2).

|  |  |
| --- | --- |
| Akron, OH | Red Wing, MN |
| Clinton, IA | Rochester, MN |
| Davis, CA | Sacramento, CA |
| Essex Junction, VT | San Jose, CA |
| Gilroy, CA | Santa Cruz, CA |
| Harrison, AR | San Mateo, CA |
| Jackson, MI | San Francisco |
| Jenison, MI | Stafford, VA |
| Kansas City, KS | Sunnyvale, CA |
| Mankato, MN | Sunnyvale, TX |
| Merced, CA | Turlock, CA |
| Napa, CA | Union Beach, NJ |
| Oceanside | Wheaton, IL |
| Orange County, FL - South | Wheeling, WV |
| Palo Alto | Wichita Falls, TX |
|  | Youngstown, OH |

### Quantification of rotavirus RNA in raw wastewater and drain samples

Raw wastewater and drainage samples (potentially containing spilled wastewater or fecal material) were collected from ten drainage channels and two wastewater treatment plants in Kampala, Uganda, from 17 and 28 March 2025. Sampling occurred over ten working days at each site, yielding a total of n=119 samples (99 drainage samples and 20 wastewater samples). Each grab sample consisted of 50 mL, treated with MgCl_2_ to achieve a final concentration of 25 mM. Samples were vacuum-filtered using S-Pak® Membrane Filters (mixed cellulose esters, pore size 0.45 μm, diameter 47 mm; Merck, Cat. No. HAWG047S6). The nucleic acids were extracted from the sample using AllPrep PowerFecal Pro DNA/RNA Kit (Qiagen, Cat. No. 80254). Filters were subsequently torn into small pieces with sterile forceps and transferred into the PowerBead Pro Tube, followed by the manufacturer’s protocol. To remove potential PCR inhibitors, 100 µL of the extracted nucleic acid was purified using the Zymo OneStep PCR Inhibitor Removal Kit (Zymo Research, Cat. No. D6030). Purified extracts were 3x diluted with nuclease-free water to minimize potential inhibition and stored at −80°C until shipment on dry ice to the Eawag Laboratory (Dübendorf, Switzerland). Upon arrival, extracts were stored at −80°C for 83 to 106 days prior to analysis. Quantification of rotavirus RNA was performed using a one-step digital RT-PCR assay on the naica® PCR system (Stilla Technologies, France). The assay employed a duplex primer/probe set for simultaneous detection of rotavirus (HEX-labeled) and norovirus GII RNA (norovirus GII not reported herein).

Each RT-dPCR reaction consisted of a total volume of 27 µL, including 5.4 µL of RNA template and 21.6 µL of mastermix. The mastermix contained qScript XLT One-Step RT-qPCR ToughMix (2x) (Quantabio, USA, Cat. No. 95132), 0.5 µM of forward and reverse primer, 0.2 µM of each probe, 0.05 µM of fluorescein sodium salt (VWR, Cat. No. 0681-100G), and RNase-free water. The sequences of the rotavirus primers and probes are: GGCTTTTAAAGCGTCTCAGT (forward), AATYTATAGCTATCRTTCTCYARATG (reverse), and HEX/CCATGGCTGAGCTAGCTTGCTT/BHQ-1 (probe)^46^.

As with the solid samples above, two different cycling conditions were used. One set of conditions did not include a heat snap and the other set included an initial denaturation step where the template was heated to 95°C for 5 min followed by cooling on ice for 5 min and then centrifuged briefly before being added to the reaction. The reaction mixture (25 μL) was loaded onto Sapphire chips (Stilla Technologies) were partitioned into droplets using a Geode system (Stilla Technologies; 12 min at 40°C), followed by thermocycling with the following conditions: reverse transcription (50°C for 1 h), enzyme activation (95°C for 5 min), and 45 cycles consisting of denaturation (95°C for 15 s) and annealing/extension (54°C for 1 min). After the reaction, the chips were scanned using a Naica Prism3 Crystal Reader (Stilla Technologies). Droplet counts and fluorescence signals across three channels (blue, green, and red) were analyzed using Crystal Miner software (Stilla Technologies). As a quality control measure, samples with < 15,000 droplets generated were considered invalid. Absolute concentration of rotavirus RNA (gc/µL) was calculated automatically by the software using Poisson distribution analysis of positive droplets, and the unit was converted into gc/mL considering the volumes of sample, extract, and PCR template and dilution factor. Samples were run in technical duplicates, along with a positive control and a no-template control (NTC). Results were expressed as copies per milliliter (gc/mL), with errors calculated as the standard deviation between duplicates. The lowest detectable concentration was 8 cp/ml.

### Ethics approval

The protocols for collecting and processing samples from Uganda were approved by the Vector Control Division Research Ethics Committee in Uganda (approval no. VCDR-2024-65) and the Eawag Ethical Review Committee.

### Statistics

The ratio of RNA measured with and without a heat snap was calculated; the ratio was found to be log-normally distributed using a Shapiro-Wilks test (p=0.18). A t-test using the log_10_-transformed data was used to test the null hypothesis that that ratio for solids is the same as for the liquid (wastewater and drainage) samples. A linear regression was used to assess the relationship between measurements made with and without a heat snap. All analyses were performed using Rstudio (version 1.4.1106) with R (version 4.0.5). All measurements are deposited in the Stanford Digital Repository (https://doi.org/10.25740/rk700qq6039).

## Results

All negative and positive controls were negative and positive, respectively indicating that assay performance was acceptable.

Twenty-three of 99 drain samples, and 3 of 38 wastewater solids samples were negative for rotavirus RNA using no heat snap, yet had detectable RNA when using a heat snap. All raw wastewater samples had detectable RNA using both approaches. These results suggest using the heat snap provides higher sensitivity than not using a heat snap.

Samples for which rotavirus RNA was detected using both with and without a heat snap are considered in the following quantitative analysis (n=131). The concentrations of rotavirus RNA measured using a heat snap were higher than those measurements without a heat snap in both wastewater solids and liquids. The median ratio was 123 (interquartile range 82 to 196, n = 131). Ratios were not different for liquids versus solids (t-test on log_10_-transformed ratios, t = 1.323, dof = 50.302, p = 0.19). The relationship between log_10_-transformed concentrations measured with and without a heat snap (C_HS_ and C_NHS_, respectively) was linear with a slope of 1.0±0.02, and intercept of 2.1±0.05 (errors represent standard deviation, r=0.98, p < 0.05), suggesting a power-law relationship: C_HS_ = 10^2.1^C_NHS_^1^. Results suggest that measurements made with a heat snap are approximately 125 times higher than those made without a heat snap.

## Discussion

Rotavirus RNA was detected consistently in samples when a heat snap was used, suggesting the heat snap increased sensitivity. In addition, higher concentrations were measured using a heat snap. This suggests that rotavirus RNA concentrations in samples based on methods that do not include a heat snap are underestimates. At least a portion of the dsRNA rotavirus genome present in wastewater solids, wastewater, and wastewater impacted drainage is not denatured endogenously. Therefore a denaturing step is needed to more accurately characterize rotavirus RNA concentrations in these matrices.

The linear relationship between the measurements with and without a heat snap is striking. We note that the relationship was conserved despite samples that were collected from diverse locations and times, and represent different matrices, and that the samples were processed in different laboratories using different extraction methods and quantified using different digital PCR methods. The linear relationship suggests that the proportion of rotavirus RNA accessible to the reverse transcription without heat snap relative to the proportion accessible after heat snap is conserved. In our analyses, the ratio is 1:125, suggesting that an average 0.8% of the rotavirus RNA is quantifiable without a heat snap. The conservation of the linear relationship in our samples also suggests that measurements made without a heat snap may be corrected to approximate those with a heat snap by multiplying by 125, though the applicability of this correction factor to other sample matrices beyond those tested here or using alternative extraction methods is uncertain.

A heat snap is likely needed prior to reverse transcription when dsRNA is not denatured endogenously in a sample or otherwise during extraction. Although the work here highlights the impact of heat snap on quantifying rotavirus, there are a number of other important human and animal dsRNA pathogens, including other members of the *Sedoreoviridae* family (i.e., Bluetongue virus and Colorado tick fever virus) and members of the *Birnaviridae* family (i.e., Infectious Bursal Disease Virus). Failure to denature dsRNA prior to reverse transcription will likely influence other studies beyond PCR-based quantification assays, such as metagenomic sequencing; dsRNA viruses may also be underrepresented in metagenomic data sets. It is unclear under what conditions a heat snap is needed, but given the results reported herein, we recommend that a heat snap be used when quantifying dsRNA from environmental matrices. We further recommend manufacturers of molecular biological products including RT highlight the need to denature dsRNA prior to RT.

## Acknowledgements

We acknowledge Katja Herter for noting this phenomenon in her M.Sc. thesis.

## Conflicts of interest

BS and DD are employees of Verily Life Sciences, LLC.

**Figure 1.**
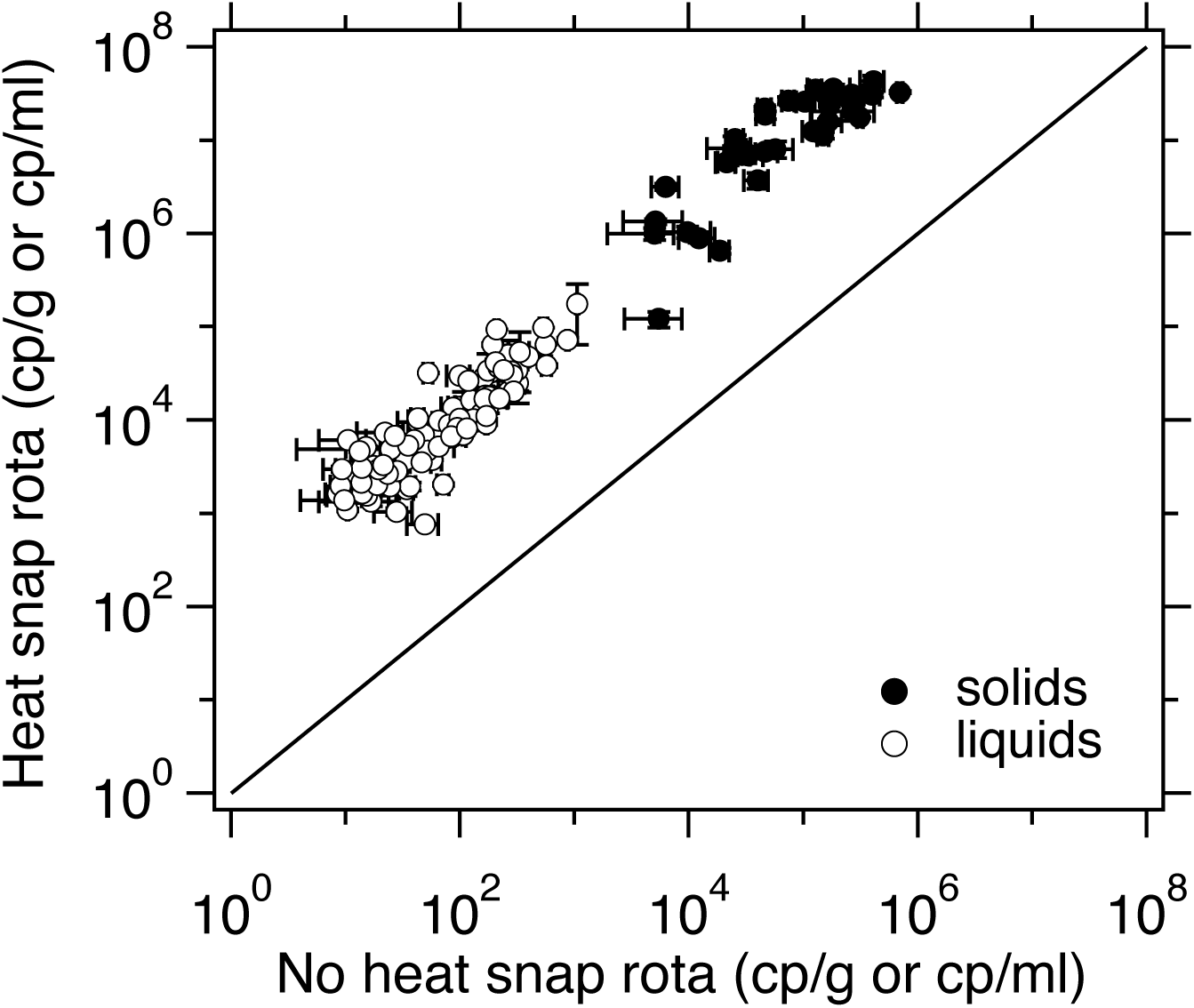
Concentrations of rotavirus RNA measured with (y-axis) and without (x-axis) a heat snap. Error bars are standard deviations; both y and x error bars are shown and if they cannot be seen then they are smaller than the symbol. Black symbols are wastewater solids, white are raw wastewater or drainage samples. Only samples where both measurements were above the lowest detectable concentration are shown. The line represents the 1:1 line. Omitted are n = 3 wastewater solid samples and n = 23 drainage samples where the sample run without a heat snap where below the lowest detectable concentration.

